# Hypothalamic neuronal activation in primates drives naturalistic goal-directed eating behavior

**DOI:** 10.1101/2023.06.11.544288

**Authors:** Leslie Jaesun Ha, Hyeon-Gu Yeo, Yu Gyeong Kim, Inhyeok Baek, Eunha Baeg, Young Hee Lee, Jinyoung Won, Yunkyo Jung, Junghyung Park, Chang-Yeop Jeon, Keonwoo Kim, Jisun Min, Youngkyu Song, Jeong-Heon Park, Kyung Rok Nam, Sangkyu Son, Seng Bum Michael Yoo, Sung-hyun Park, Won Seok Choi, Kyung Seob Lim, Jae Yong Choi, Jee-Hyun Cho, Youngjeon Lee, Hyung Jin Choi

## Abstract

Eating addiction is the primary cause of modern obesity. Although the causal role of the lateral hypothalamic area (LHA) for eating is demonstrated in rodents, there is no evidence in primates regarding naturalistic eating behaviors. We investigated the role of LHA GABAergic (LHA^GABA^) neurons in eating by chemogenetics in three macaques. LHA^GABA^ neuron activation significantly increased naturalistic goal-directed behaviors and food motivation, which was specific for palatable food. PET and MRS validated the chemogenetic activation. Rs-fMRI result revealed that functional connectivity (FC) between the LHA and frontal areas was increased, while the FC between the frontal cortices was decreased after the LHA^GABA^ neuron activation. Thus, our study elucidates the role of LHA^GABA^ neurons in eating and obesity therapeutics for primates and humans.

## Introduction

Eating addiction and impulsive goal-directed eating behaviors is the major cause driving the rapid rise of obesity and related contemporary health problem worldwide *(1)*. Hypothalamus is the major regulator of eating. However, the literature linking the hypothalamus to goal-directed eating behaviors are limited to rodents. No study clearly demonstrated the role of the hypothalamus on naturalistic goal-directed eating behavior for human and non-human primates.

Within the hypothalamus, the lateral hypothalamic area (LHA) has been investigated as the critical regulator of eating and eating-related goal-directed behaviors for over 70 years in all animals, including humans *(2-5)*. We and others have demonstrated that among the diverse cell types in the LHA, GABAergic (LHA^GABA^) neurons are the crucial candidate for food motivation and goal-directed behaviors *(6-10)*. In addition, LHA^GABA^ neuron activation in rodents increased aberrant gnawing behavior *(11-13)*. However, due to differences between species, research on rodents is not directly translated into humans. Non-human primate studies, on the other hand, can bridge the gap between rodent and human studies, as macaques genetically closest to humans exhibit higher cognitive intelligence for goal-directed behavior compared to rodents *(14)*. Nevertheless, the role of LHA^GABA^ neurons in non-human primates is still unknown.

## Results

### Experimental scheme of LHA^GABA^ neuron activation by chemogenetics and confirmation of viral injection and expression

The general experimental scheme is shown in Fig. 1A. Three female rhesus macaques (Monkeys A, B, and C) were injected with an adeno-associated virus (AAV) of AAV9-hDlx-hM3Dq-dTomato in the LHA, which induces GqDREADD nuclear dTomato expression in GABAergic interneurons under the GABA neuron-specific promoter (hDlx enhancer). The experimental tasks were conducted seven weeks after the virus injection. The monkeys were injected with clozapine N-oxide (CNO in 10% dimethyl sulfoxide, DMSO) for LHA^GABA^ neuron activation or the vehicle (saline in 10% DMSO) for the control. Multiple cameras were used to observe the behavior of the monkeys with various experimental stimuli: palatable food, unpalatable food, water, and non-food object. The analytical methods were classified into manual and deep learning-based for the sequential goal-directed behaviors. For the manual analysis, behavioral indices of ‘approach hands to goal,’ ‘bring the goal to the mouth,’ and ‘food peeling’ were assessed. For the deep learning-based analysis, the indices of ‘tray approach,’ ‘bout,’ and ‘duration in food zone’ were assessed. The classification of behavior phases and the behavioral index are presented in Table S1.

**Fig. 1.**
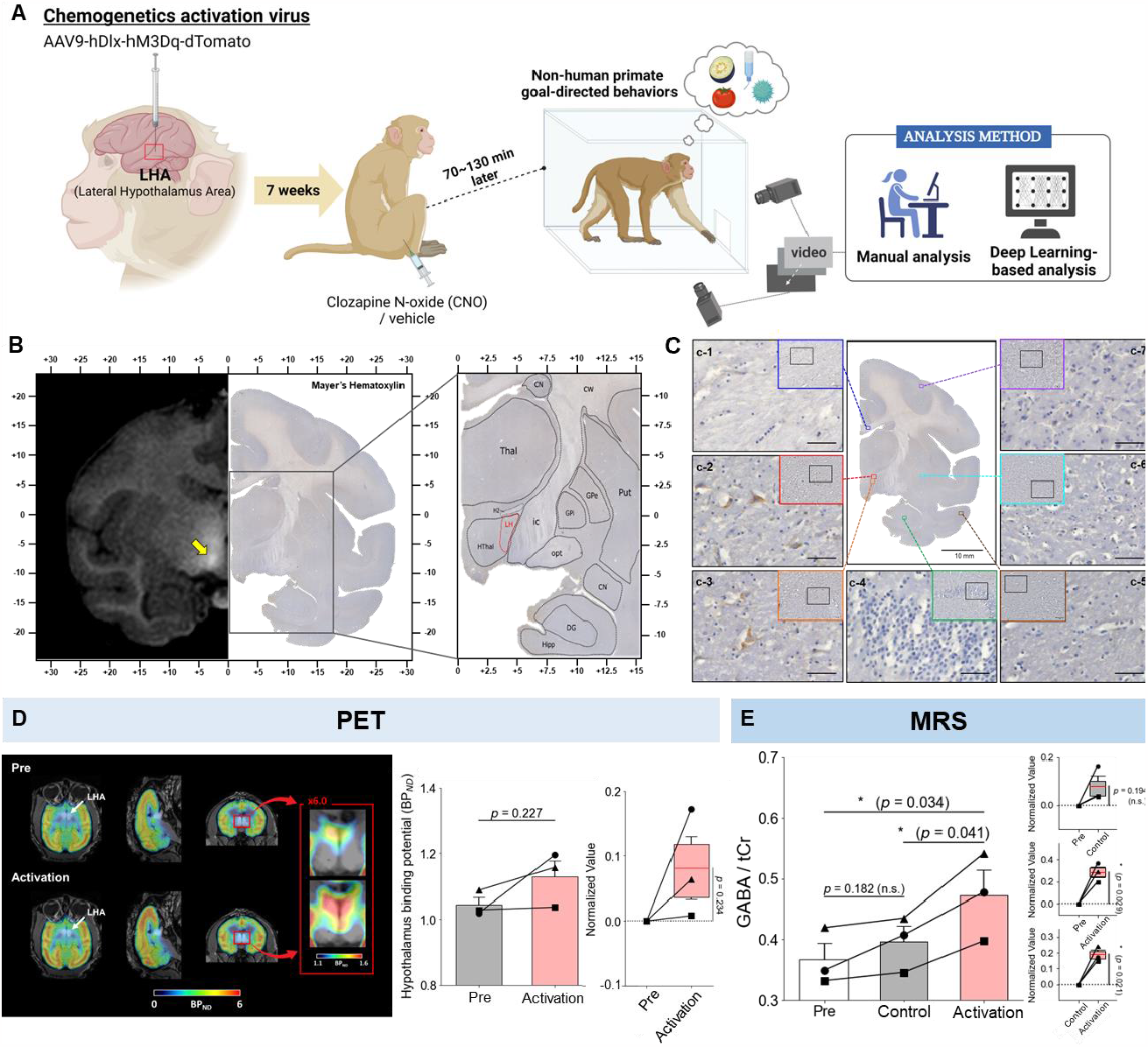
Experimental scheme of LHA^GABA^ neuron activation via chemogenetic and virus validation by histological and functional imaging. (**A**) Schematic of viral injection for LHA^GABA^ activation. After the virus expression, the monkeys were injected by CNO as the LHA^GABA^ activation and vehicle as the control. During the peak chemogenetic effect time window, the behavior of monkeys was simultaneously observed by multiple cameras with various experimental stimuli (palatable food, unpalatable food, water, and non-food object) and analyzed by manual and deep learning based. (**B**) Confirmation of viral injection by MRI and histology (AAV injection area). (left) MRI image after contrast agent injection co-infused with AAVs and histological image of matching region. The yellow arrow indicated the LHA region. (right) Enlarged landmark for injection area. Black dotted line indicates brain subregions and red dotted line indicate AAV injection site. CN; caudate nucleus, cw; cerebral white matter, DG; dentate gyrus, Gpe; globus pallidus (external division), Gpi; globus pallidus (internal division), H2; field H2 (of Forel), Hipp; hippocampus, Hthal; hypothalamus, ic; internal capsule, opt; optic tract, Put; putamen, Thal; thalamus. (**C**) Confirmation of viral expression by immunohistochemistry. Yellow arrows point to RFP-labeled neurons in LHA. c-1; corpus callosum, c-2; LHA (superior), c-3; LHA (inferior), c-4; hippocampus, c-5; cortex (temporal lobe), c-6; putamen, c-7; cortex (parietal lobe). Scale bar: 50 μm. (**D**) Comparison of GABA PET-MR images on Monkey A. (left) [^18^F]flumazenil binding in the non-human primate at pre and after administration of CNO as an activation were fused with MR. White arrow indicates lateral hypothalamus and red arrow represents magnification images of hypothalamus (x6.0). (right) Comparison of hypothalamus binding potential value at pre and activation (compared by paired t-test). (**E**) The comparison of pre, control and activation session on Monkey A, B, and C (compared by paired t-test). Throughout the whole figure, the circle, square, and triangle markers correspond to Monkey A, B, and C, respectively. The mean value of GABA/tCr and normalized value were presented. Those in the right side of (D, E) show the rate of change caused by control or activation compared to pre or control injection (compared with 0 by one-sample t-test). *p*-value; * p < 0.05.

### Confirmation of viral injection site and histological validation of the virus expression by immunohistochemical staining (IHC)

We injected the virus into the LHA using gadolinium contrast magnetic resonance imaging (MRI) confirmation. LHA targeting was further confirmed using histologic slides stained with hematoxylin and based on landmarks established *(15)* (Fig. 1B). Immunohistochemical staining (IHC) was performed on one subject (Monkey A) with anti-red fluorescent protein (anti-RFP) antibody to confirm AAV-mediated chemogenetically induced gene expression in the LHA to validate the viral expression at the site of the LHA (Fig. 1C). The results showed that RFP-labeled neurons were detected only in the LHA, and no immunoreactivity was observed in other regions, such as the corpus callosum, hippocampus, temporal cortex, putamen, and parietal cortex. An additional subject (Monkey D) was injected with the virus in the LHA to further validate the virus’s performance (fig. S1). From the anterior to the posterior (fig. S1A), the immunohistochemistry result showed that RFP-labeled neurons were detected only in the LHA (fig. S1, B to D, arrows) and not in other areas. The results confirmed that AAV-mediated chemogenetically induced gene expression was only present in the targeted brain region.

### Functional validation of the GABA system using in vivo positron emission tomography (PET) imaging and proton magnetic resonance spectroscopy (^1^H-MRS)

We performed GABA PET with a specific PET radiotracer for GABA-A receptors (i.e., [^18^F] flumazenil) to quantify and validate LHA^GABA^ neuron activation *in vivo*. Through MR-guided PET experiments, we confirmed LHA^GABA^ neuron activation in the bilateral hypothalamus (Fig. 1D, figs. S2, A and B, white arrow). After LHA^GABA^ neuron activation, binding potential values were consistently increased in the hypothalamus for all three monkeys (Fig. 1D). In addition to the hypothalamus, increases in the binding values were observed in other areas (figs. S2, C to E).

We performed GABA MRS analyses to quantify the changes in the metabolite concentrations to further validate the *in vivo* imaging of GABA concentration. A 7T MR scanner was used to measure the GABA/tCr value normalized as a ratio of GABA to total creatine (creatine + phosphocreatine, tCr) (Fig. 1E). The brain anatomical reference and the fitting data obtained by the semi-localized adiabatic selective refocusing (sLASER) method are shown in fig. S3, A and B. An individual representative result of real-time dynamics changes in GABA/tCr is shown in fig. S3C. LHA^GABA^ neuron activation significantly increased the GABA/tCr value in the LHA (Fig. 1E and fig. S3D).

The GABA system change after LHA^GABA^ neuron activation for individual monkeys showed a significant positive correlation between PET and MRS (fig. S4A). Further, the monkey with a higher effect size on PET also showed a higher effect size on MRS (Monkey A > Monkey C > Monkey B) (fig. S4, B and C). These results indicate that the effect size of LHA^GABA^ neuron activation can be quantified through PET and MRS. Collectively, these functional results confirm the chemogenetic activation of LHA^GABA^ neurons.

### Goal-directed eating behavior for palatable food was significantly increased after LHA^GABA^ neuron activation

We quantified goal-directed eating behaviors for palatable food with or without the chemogenetic activation of LHA^GABA^ neurons to provide efficacy of naturalistic behavioral outcomes. Behavior temporal dynamics analysis showed that LHA^GABA^ neuron activation increased the time spent in the food zone with the food consumption posture (Figs. 2, A and B). The raster plot of Fig. 2B shows that activating LHA^GABA^ neurons increased the duration of low locomotion and goal-directed behavior indices, compared to the control in Fig. 2A. We manually annotated three behavioral indices to quantify naturalistic goal-directed behavior: approaching hands to food, bringing the food to the mouth, and food peeling (Table S1). The number of approaching hands to food, bringing the food to the mouth, and food peeling behavior was significantly increased after LHA^GABA^ neuron activation (Figs. 2, C to E). Further, food intake was consistently increased in all three monkeys after LHA^GABA^ neuron activation (Fig. 2F). The total duration of low locomotion was significantly increased after LHA^GABA^ neuron activation (fig. S6A).

**Fig. 2.**
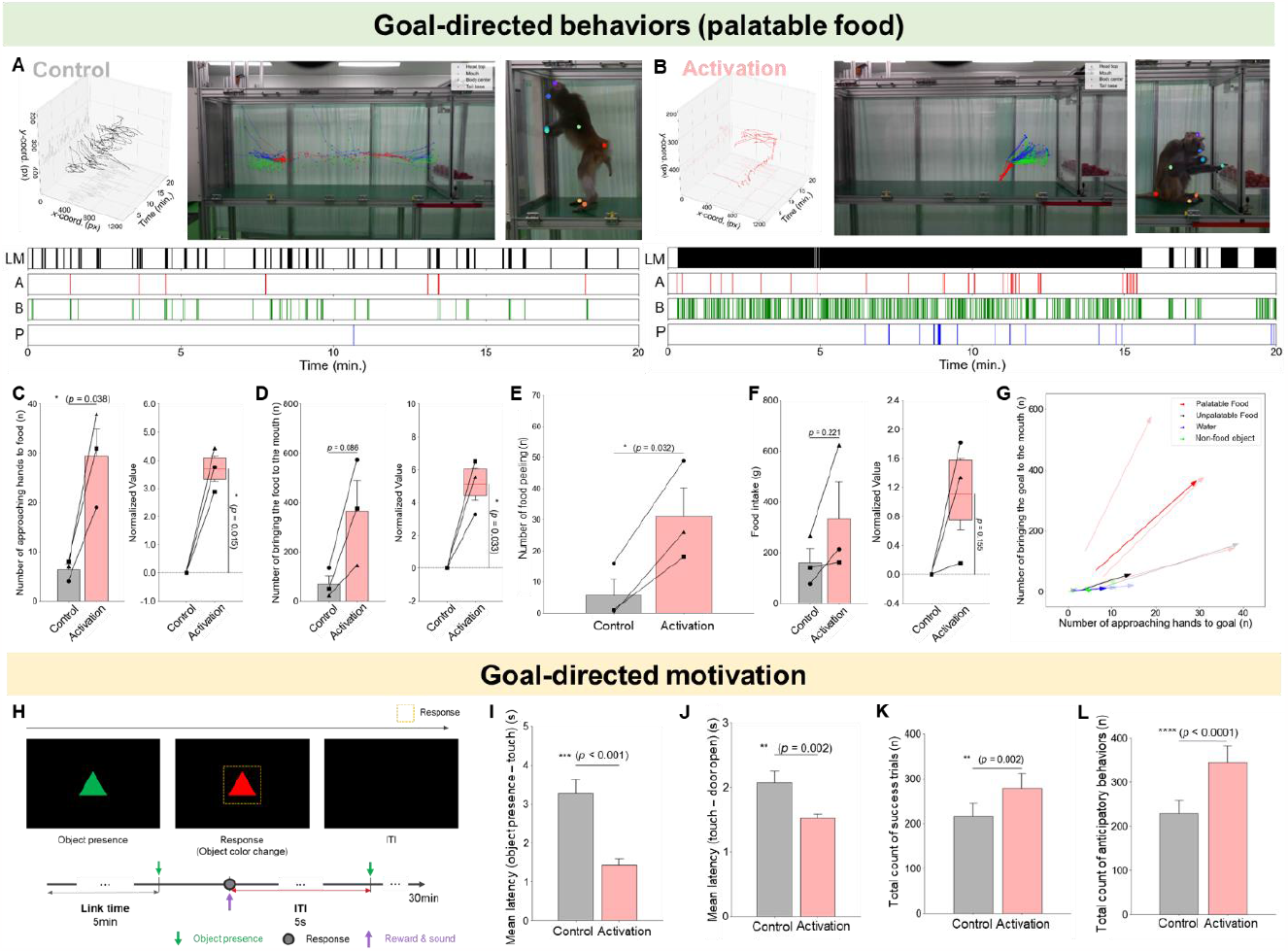
Goal-directed eating behavior and motivation for palatable food was significantly increased after the LHA^GABA^ neuron activation. (**A**) Behavior temporal dynamics analysis of control. (top) The 3D trajectory of body center (average over an interval time window of 15 sec. length) and the posture of Monkey B (first 5 min.). (bottom) The raster plot of LM; low locomotion duration, A; approach hands to food, B; bring the food to the mouth, and P; food peeling. (**B**) Behavior temporal dynamics analysis of LHA^GABA^ neuron activation. (**C**) The number of approaching hands to food. (**D**) The number of brining the food to the mouth. (**E**) The number of food peeling. (**F**) The intake of palatable food. (**G**) The 2D plotting of number of approaching hands to goal and number of bringing the goal to the mouse on all experimental stimuli (red arrow; palatable food, black arrow; unpalatable food, blue arrow; water, green arrow; non-food object). (**H**) The schematic of operant conditioning paradigm on FR1. (**I**) Mean latency over all trials between control and LHA^GABA^ neuron activation. The absolute *p*-value was 4.52e^-7^. (**J**) Mean latency over all trials between control and LHA^GABA^ neuron activation. (**K**) Total count of success trials. The error bars indicate 95% confidence intervals from bootstrapping. The absolute *p*-value was 0.002. (**L**) Total count of anticipatory behaviors. The error bars indicate 95% confidence intervals from bootstrapping. The absolute *p*-value was 6.479e^-7^. The figures in the left side show the values of corresponding indices in Monkey A, B, and C between control and LHA^GABA^ neuron activation (compared by paired t-test). Those in the right side show the rate of change caused by activation (compared with 0 by one-sample t-test). Throughout the whole figure, the circle, square, and triangle markers correspond to Monkey A, B, and C, respectively. The red line under the cage of picture indicates the food zone. *p*-value; * p < 0.05, ** p < 0.01, *** p < 0.001, **** p < 0.0001.

The effect of LHA^GABA^ neuron activation was specific to palatable food stimuli. Contrary to palatable food, no definite efficacy was observed for unpalatable food after LHA^GABA^ neuron activation (figs. S5, A to F). For unpalatable food, there were no significant changes in the food consumption posture, the time duration in the food zone, and goal-directed eating behavioral indices after LHA^GABA^ neuron activation (figs. S5, A to F). Further, to find whether LHA^GABA^ neuron activation influences goal-directed behavior for other context stimuli, water (fig. S5G) and a non-food object (fig. S5K) were tested. For both water and the non-food object, no significant changes in goal-directed behaviors were observed after LHA^GABA^ neuron activation (figs. S5, G to M). Further, there were no significant change in the total duration of low locomotion for unpalatable food, water, and non-food object after the LHA^GABA^ neuron activation (figs. S6, B to D).

Collectively, we visualized a 2D plot to elucidate the effect of LHA^GABA^ neuron activation on two main goal-directed behavioral indices: “approach hands to the goal” and “bring the goal to the mouth” (Fig. 2G) for four different context stimuli, which included, palatable food, unpalatable food, water, and a non-food object. The results demonstrated profound changes only for the palatable food stimuli. However, no robust effects of LHA^GABA^ neuron activation were observed for other stimuli (unpalatable, water, and non-food object).

In addition to a manual analysis, deep learning-based analysis also showed that the goal-directed eating behaviors were consistently increased only for palatable food after LHA^GABA^ neuron activation. We conducted a deep learning-based analysis for the palatable food and unpalatable food throughout the experiment (figs. S7, A to H). LHA^GABA^ neuron activation consistently increased the number of tray approaches, the number of bouts, and the duration in the food zone in all three monkeys (figs. S7, A to D). The deep learning-based analysis of hand-to-tray distance, bout, and food zone duration showed a higher frequency and temporal density and a closer distance in the goal-directed eating behaviors after LHA^GABA^ neuron activation (fig. S8). However, no robust change in behavioral indices was observed for unpalatable food (figs. S7, E to H).

In summary, goal-directed eating behavior for palatable food was significantly increased after LHA^GABA^ neuron activation (movie S1). No definite change in goal-directed behavior was observed for unpalatable food, water, and the non-food object after LHA^GABA^ neuron activation (Table S1). In addition, contrary to that in rodent studies (*11-13*), no aberrant gnawing behavior was observed with both LHA^GABA^ neuron activation and the control for all stimuli: palatable food, unpalatable food, water, and non-food object.

### Goal-directed motivation for palatable food was increased after LHA^GABA^ neuron activation

We conducted an operant conditioning paradigm using a touchscreen to investigate whether LHA^GABA^ neuron activation increases motivation for palatable food. A sweet pellet was delivered every time the monkey touched a screen object (FR1 experimental scheme, Fig. 2H; movie S2 and S3). The mean latency of the object-to-touch time and touch-to-door open time was significantly shorter after LHA^GABA^ neuron activation than that of the control (Fig. 2, I and J; fig. S9, A and B), indicating an increased desire to eat. The number of successful trials was significantly higher for LHA^GABA^ neuron activation (Fig. 2K and fig. S9C). The anticipatory behavior was quantified by the number of magazine door touching instances during the no-reward period. The number of anticipatory behaviors was significantly higher with LHA^GABA^ neuron activation (Fig. 2L and fig. S9D). These results demonstrate that the activation of LHA^GABA^ neurons increases the motivation and the desire to eat palatable food.

### Functional connectivity (FC) changes after LHA^GABA^ neuron activation measured by resting-state functional MRI (rs-fMRI)

We measured pre-scan and post-scan rs-fMRI before and after the LHA^GABA^ neuron activation or control (Fig. 3A) with 198 parcellated regions of interest (ROIs) (Fig. 3B) in two monkeys to investigate the effect of LHA^GABA^ neuron activation on FC. The LHA seed-based FC analysis revealed a significant increase in FC, primarily between the frontal areas and the LHA, which are functionally and structurally connected (Figs. 3, C and E; and fig. S10B). Whole-brain network analysis revealed a significant decrease in FC, primarily within the frontal cortices after LHA^GABA^ neuron activation (Figs. 3, D and F; and fig. S10C). The results indicate that the FC between the LHA and frontal areas is increased, while the FC between frontal cortices is decreased after LHA^GABA^ neuron activation.

**Fig. 3.**
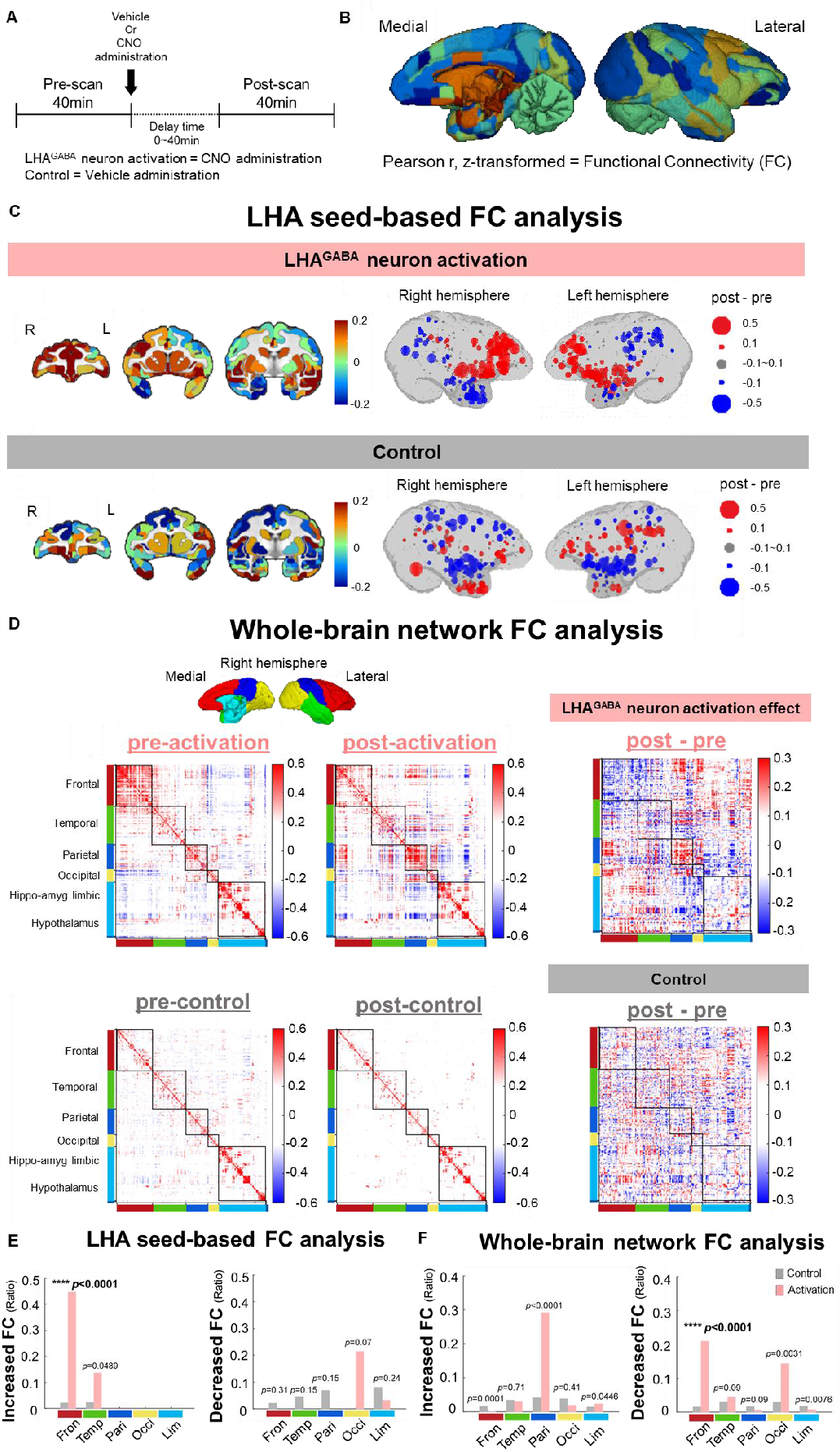
Functional connectivity (FC) change after the LHA^GABA^ neuron activation measured by resting-state functional MRI (rs-fMRI). (**A**) Time course of acquiring cerebral blood volume (CBV)-weighted rs-fMRI scans using MION, a T2 contrast. (**B**) 198 parcellated ROIs to calculate FC using the segmentation map of the D99 atlas. (**C**) LHA seed-based FC analysis. FC differences between LHA and other ROIs. (left) 2D coronal section ROI visualization. (right) 3D whole brain ROI visualization. (**D**) Whole-brain network FC analysis. (left) FC matrices between ROIs during pre-activation/control, (middle) post-activation/control, and (right, LHA^GABA^ neuron activation effect/control) the difference. FCs were sorted in a lobe-dependent manner, as shown in the upper colored medial and lateral brain images. (**E**) Proportion of significantly increased and decreased FCs in LHA seed-based FC analysis. The absolute *p*-value of frontal cortex region in increased FC was 1.1042e^-6^. (**F**) Proportion of significantly increased and decreased FCs in whole-brain network FC analysis. The absolute *p*-value of frontal cortex region in decreased FC was 2.3512e^-48^. The whole-brain network FC changes used the matrices of the black rectangles in (D) and other methods were same as in (E). The FC changes were considered significant if they deviated from the 95% threshold of the control FC distribution, which is constructed with FC changes of control (vehicle). A chi-square test was used to compare the frequencies of FC changes. *p*-value; **** p < 0.0001.

## Discussion

Here, we investigate the role of LHA^GABA^ neuronal effects on various aspects of naturalistic goal-directed behavior in freely moving non-human primate via chemogenetic technique. Our study demonstrated that LHA^GABA^ neuron activation in non-human primates drives only palatable food-specific goal-directed eating behavior.

LHA^GABA^ activation in non-human primate, unlike in rodent studies, evoked naturalistic goal-directed eating behavior only for palatable food. We and others reported that the activation of LHA^GABA^ neurons increases food motivation in rodents *(6-10)*. In addition, other studies in rodents reported that these neurons mediate aberrant gnawing behavior regardless of the calorie content, including for wood and water *(12, 13)*. However, interestingly, according to our results, LHA^GABA^ neuron activation in non-human primates increased the goal-directed eating behavior and food motivation only for the palatable food stimuli (Fig. 2). Specifically, after the LHA^GABA^ neuron activation, the monkeys had decreased locomotion to focus on eating behaviors, only for the palatable food. They displayed sequential naturalistic eating behaviors such as approaching, peeling, and bringing the food to the mouth without aberrant gnawing behavior. Furthermore, after the LHA^GABA^ neuron activation, increased motivation for the palatable food was observed during the operant conditioning paradigm. Regarding the mechanistic difference between rodents and macaques, we speculate that LHA^GABA^ neurons in non-human primates may have different cell type characteristics, anatomical distribution, or projection circuitry compared to those in rodents *(7-10)*. Indeed, we recently reported a food-specific subpopulation of LHA^GABA^ neurons in rodents *(10)*. In addition, interestingly, LHA deep brain stimulation successfully treated refractory obesity patients *(4)*. In this regard, the present study suggests promising therapeutic strategies targeting specific cell types in LHA for human obesity.

The present rs-fMRI results provide further mechanistic evidence highlighting the role of the frontal-LHA circuitry in regulating food-related motivation and behavior (Fig. 3). The frontal cortex is closely connected with the LHA *(16)*. Therefore, as expected, the frontal cortex exhibited the most robust change in FC in response to LHA^GABA^ neuron activation. Since the prefrontal cortex provides the major top-down inhibitory control to impulsive goal-directed motivation, the decreased intra-frontal cortex FC could be the mechanistic link contributing to the observed increased goal-directed motivation after LHA^GABA^ neuron activation.

Our PET and MRS finding validate the biological efficacy of chemogenetics after LHA^GABA^ neuron activation. In addition, increased GABA binding values were observed in the overall cortices and LHA. We speculate that these are the secondary effects of the downstream regions of LHA^GABA^ neurons (e.g., hippocampus, insula, cortex). Further, we assumed that the neuronal inhibition by increased GABA concentrations on PET could be related to the decreased inter/intra-frontal cortex FC in rs-fMRI results.

Our study of three macaques demonstrates the preclinical efficacy of naturalistic goal-directed eating behavior, which is most comparable to the human clinical context.

Chemogenetics is a new gene therapy technology for patients that targets an engineered receptor to specific cell types involved in nervous system dysfunction, allowing for highly selective drug-controlled neuromodulation (*17*). A non-human primate models study demonstrated the feasibility of the chemogenetics for the treatment of humans (*18*). In recent years, chemogenetics in primates has been widely used to target specific neuronal populations in awake states. However, these chemogenetic studies were restricted to non-naturalistic or non-goal-directed behaviors. Most studies used head-fixed primate chairs to investigate the reward value *(19-22)* and saccade *(21, 23-26)*. The other studies were non-naturalistic with non-locomotive setting (hand movement *(27-30)*, memory task *(31-35)*, social choice task *(36)*) or non-goal-directed behaviors (freezing and anxiety behavior *(18, 37, 38)*, general locomotion *(39)*, sleep efficiency *(40)*, and rotation behavior *(41)*).

Therefore, the present study provides the most naturalistic and human-like preclinical proof-of-concept evidence of inducing goal-directed eating behavior by chemogenetically activating LHA^GABA^ neurons in three macaques.

In conclusion, this study elucidates the role of LHA^GABA^ neurons in eating, increasing naturalistic goal-directed behavior toward palatable food via chemogenetic activation in non-human primates. These results open groundbreaking breakthroughs for the neurobiology of obesity and the development of obesity therapeutic strategies including drug, device, and drug-controlled gene therapy (chemogenetics).

## Materials and Methods

### Experimental animals

Three female rhesus monkeys (*Macaca mulatta*) aged 7–8 years, at 5–6 kg body weight (monkeys A, B, and C), were selected with similar temperament based on temperament assessments for the behavioral task *(42)*. For the pilot study of virus validation, the other rhesus monkey D was used. The monkeys were obtained from Guangzhou Monkey King Biotechnology Co. Ltd. (Guangdong Province, China). They were maintained in indoor cages at the National Primate Research Center of the Korea Research Institute of Bioscience and Biotechnology (KRIBB). They were fed commercial monkey chow (Teklad 2050™, Envigo, USA) and a daily supplement of various fruits and water *ad libitum* under controlled environmental conditions, including a temperature of 24 ± 2 °C, relative humidity of 50 ± 5%, and a 12:12-hour light-dark cycle. The attending veterinarian monitored the monkeys’ health according to the recommendations of Weatherall et al.’s research *(43)*. Animal health care, including microbiological tests for B virus, simian retrovirus, simian immunodeficiency virus, simian virus 40, and simian T-cell lymphotropic virus, were periodically monitored and managed. All procedures were approved by the Korea Research Institute of Bioscience and Biotechnology Institutional Animal Care and Use Committee (Approval No. KRIBB-AEC-21319) and complied with the Animal Research Reporting of In Vivo Experiments (ARRIVE) guidelines *(44)* and National Institutes of Health guidelines in the USA.

### General training procedures

All behavioral task monkeys were acclimatized to the designed experimental cage and food for 30 min before the main experiment so they could participate harmoniously. The main experiment was conducted for 20 min during the peak chemogenetic effect time window of 70 to 130 min after injection. The monkeys were administered intramuscular (i.m.) CNO (10 mg/kg) or vehicle in 10% dimethyl sulfoxide (DMSO) injection. All monkeys executed the conditioning experiment at least once a week to increase endurance to the pain they would suffer when subjected to an i.m. CNO or vehicle administration. Before and after each main experiment, the cage was disinfected with 70% alcohol. The total duration of the experiment was 180 days with twice-weekly tasks (each on a separate day).

Further, we executed four experimental tasks as follows; 1. palatable food task; cherry tomatoes were given 2 kg, which was known as a preferred food for non-human primates. The monkeys had free access and enough quantity to eat for experimental time. 2. unpalatable food task; eggplants were provided 1∼2 kg, which was less preferred by them. The monkeys had free access and had enough quantity to consume during experimental time. 3. water task; all monkeys were denied access to water for 24 hours before the experiment. Tap water at room temperature in a previously sanitized 1 L bottle was provided at the top middle of the experimental cage. The monkeys could access freely and drink without restriction or effort during the experimental time. 4. non-food object task; a lighting ball was provided that illuminated when the monkeys chewed during the experimental time.

Three cameras (HDR-CX405, SONY, Japan) for the analysis were set on top, front, and at the side of the chamber, along with a white panel equipped adjacent to the tunnel entrance to obscure the outside vision. After handling the image of the video, we analyzed the data based on manual and deep learning. The cage comprises a custom-made translucent plexiglass tunnel (195 * 51 * 84 (cm)) with a feeding tray inside. An entryway enabled an individual primate transfer cage to be docked in so monkeys could voluntarily step into the experimental cage using a sliding door.

### Surgical procedures

We injected hDlx promoter-containing AAV, AAV9-hDlx-GqDREADD-dTomato (Addgene viral prep # 83897-AAV9), into the LHA of four rhesus monkeys to achieve GABAergic neuron-specific chemogenetic gene expression. Animals were initially anesthetized via the intraperitoneal injection of a cocktail mixture of ketamine (5 mg/kg) and atropine (0.02 mg/kg) and fixed in the prone position using a custom-built CT and MRI-compatible stereotaxic frame under isoflurane-induced anesthesia (1.5% in 2 L/min oxygen). After confirmation of the head-restrained animal within the stereotaxic frame, the skin was gently separated, and the skull was widely exposed. The fiducial MRI marker was attached to the skull surface as a precise reference point. Then, a brain T1-weighted MRI was performed as a baseline reference. Baseline images were used to determine the stereotaxic coordinates of a targeted brain region. Two circular craniotomies centered on the skull surface towards a targeted coordinate were performed at each hemisphere, approximately 5–7 mm in diameter. A total of 30 μL AAV was injected using the Hamilton syringe and micro-infusion pump by convection-enhanced delivery into the LHA at each hemisphere. An initial infusion rate of 0.2 μL/min was applied and increased sequentially to 0.4, 0.6, and 0.8 μL/min at 5 min intervals. A gadolinium-based MRI contrast agent was co-infused with the viral vector (1:200) to verify the correct injection site. After the infusion, the syringe needle was kept in the targeted region for 10 min before retraction and slowly removed from the brain. Vital signs, including heart rate, pO2, and body temperature, were monitored during anesthesia throughout all surgical procedures. Antibiotics (enrofloxacin) and analgesics (ketoprofen) were administered (5 mg/kg, i.m.) after the surgery.

### Immunohistochemistry

The whole brain was removed from the skull, washed with cold PBS, post-fixed in 4% PFA for 24 h at 4°C, and sliced into 8 mm coronal slices using a custom-built monkey brain slicer. Brain slices were immersed in 30% sucrose in PBS. After dehydration of brain tissue, brain slices were embedded in Optimal Cutting Temperature (OCT) compound, and the embedded brains were frozen at -80°C. The embedding blocks were cut on a cryostat into 20 μm thick sections and blocked for 30 min with 4% BSA and then incubated for 1 hour with the following primary antibodies: polyclonal rabbit anti-RFP (1:200, Abcam, Cambridge, UK).

The appropriate biotinylated anti-rabbit (1:200, Vector Laboratories) was incubated for 30 min at room temperature. Staining was visualized using the ABC method (PK-6100, Vector Laboratories, Burlingame, CA, USA) with 3, 30-diaminobenzidine (DAB) as the peroxidase substrate. All slides were counterstained with Mayer’s hematoxylin (DAKO Agilent, Santa Clara, CA, USA).

### Radiochemistry

We used [^18^F]flumazenil as a PET radiotracer to the GABA-A receptor to evaluate in vivo functional changes of the GABAergic system in the brain. The radiotracer was synthesized by performing nucleophilic substitution of F-18 on the iodonium tosylate precursor, as described by Moon *(45)*. The radiochemical purity at the end of the synthesis was higher than 99%.

### PET image analysis

A study-specific brain template was used to analyze non-human primate brain data. Each MR image was spatially normalized to the Rhesus Macaque INIA 19 brain MRI template embedded in PMOD software (version 4.3, PMOD Technologies Ltd., Switzerland). After static PET images (4–26 frames) were obtained from the dynamic PET image, each was co-registered with the raw MR image. The mean PET image was spatially normalized to the MR template. Finally, individual dynamic PET images were spatially normalized to the MR template.

The uptake value in the PET image was normalized in units of standardized uptake value multiplied by the body weight and classified by the injected dose. The decay-corrected regional time-activity curves were acquired from volumes of interest (VOIs), including the hippocampus, frontal cortex, motor cortex, occipital cortex, temporal cortex, parietal cortex, insular, somatosensory cortex, hypothalamus, and pons (fig. S2C). Non-displaceable binding potential was estimated using a parametric simplified reference tissue model with the pons as a reference region to estimate the density of available receptors *(46)*.

### MRI

MRI experiment was conducted on a 3.0-T MRI scanner (Achieva 3.0T, Philips Medical Systems, Best, The Netherlands) with an 8-channel Knee coil. Three-dimensional (3D) sagittal T1-weighted images were acquired using Turbo Field Echo sequence, with the following parameters: TR/TE = 14/6.8 ms, field-of-view (FOV) = 128 × 128 mm2), matrix size = 256 × 256, acquisition voxel size = 0.5 × 0.5 × 0.5 mm3 and the number of slices = 150, average = 4. Details of MRI protocols were the same as in the previous report *(47)*.

### MRS experiments

The monkeys were initially anesthetized via the intraperitoneal injection of a cocktail mixture of ketamine (0.5 mg/kg) and atropine (0.02 mg/kg). During the MRI scan, respiratory anesthesia was maintained with 2.0–2.5% isoflurane and pure oxygen (2 L/min) through a mask in the supine position. MRI experiments were performed on a 7T whole-body MR scanner (Achieva 7.0T, Philips Medical Systems, Best, The Netherlands) equipped with a 32-channel receive head coil and a birdcage 8-channel transmit coil (Nova Medical) at the Korea Basic Science Institute. For brain anatomical reference, as shown in fig. S3A, a three-dimensional magnetization prepared rapid gradient echo (3D MPRAGE) sequence was employed with the following parameters: field of view = 150 × 150 × 150 mm^3^, voxel size = 0.9 × 0.9 × 0.9 mm^3^, slices = 167, repetition time (TR) = 5.5 ms, echo time (TE) = 2.6 ms, inversion delay time = 1,300 ms, flip angle = 7°, SENSE factor = 2). Single voxel ^1^H-MRS was performed to measure the change of GABA concentration in LHA before and after the injection of CNO (10 mg/kg) in 10% DMSO. A VOI was measured by the size of 25 × 20 × 15 mm^3^ in the LHA (Fig. 2C, yellow box). The sequence and parameters are as follows: semi-localized by adiabatic selective refocusing (sLASER), TR = 5,000 ms, TE = 33 ms, spectral bandwidth = 4 kHz, samples = 2,048, average = 64, acquisition time = 5 m 30 s.

A total of 20 identical MRS scans were performed consecutively: 3 times before vehicle injection as a pre-control, 7 times after vehicle injection as a control to see the effect of vehicle only, and 10 times after CNO injection to see the effect of LHA^GABA^ neuron activation. In all spectral acquisition, variable pulse power and optimized relaxation delays (VAPOR) were used for water suppression. Before ^1^H spectra collection, the second order B_0_ shimming by the Philips pencil beam volume algorithm and RF power optimization on the localized VOI were applied. After the MRS scans, the brain anatomical reference image was reacquired to confirm that the monkey did not move during the MRS scans.

### MRS data processing

All MRS data were fitted using the LCModel software, and the GABA concentrations were normalized as a ratio of total creatine (creatine + phosphocreatine, tCr). The basis set used in LCModel was simulated in the Versatile Simulation, Pulses, and Analysis (VeSPA) package program, and LCModel provided the macromolecule basis set. Cramer-Rao lower bounds (CRLB) were used to evaluate the reliability of the fitting, and GABA metabolites with CRLB < 25% were included for further analysis. A representative example of LCModel fit results for the MR spectrum is shown in fig. S3B. The total and GABA spectra were successfully fitted, showing a good spectral dispersion with high SNR.

### Resting-state functional MRI (rs-fMRI) experiments

To investigate the change of functional connectivity (FC), we employed rs-fMRI (Achieva 3.0T, Philips Medical Systems, Best, The Netherlands) in two monkeys. A T2 contrast agent (MION (BioPAL Inc., Worcester, MA, US), 10 mg/kg) was administered before the 80∼120 min of data acquisition to improve the sensitivity and specificity of the images. A dose of ketamine (5 mg/kg) was initially administered to the monkeys, after which they were intubated, placed in an MR-compatible stereotaxic apparatus, and maintained under 1%–2% isoflurane. Scanning included the acquisition of images for 40 min (8 runs) functional scan sequence during pre- and post-activation or control (TR = 2 s, TE = 25 ms, spatial resolution = 1 × 1 × 1.5 mm^3^, FOV = 128 × 128 × 45 mm^3^, 104 vol (3.5 min)).

### Resting-state functional MRI (rs-fMRI) image processing and data analysis

Data preprocessing and subsequent analyses were primarily carried out using FSL (FMRIB’s Software Library) and MATLAB. The structural T1-weighted image was co-registered with the mean functional image and manually oriented to the D99 template *(48)*. We used 198 parcellated regions of interest (ROIs) for the measurements. Using the resulting parameters, we rotated the functional image accordingly. We then applied several steps to the fMRI data, including skull stripping, motion correction, normalization, bandpass filtering (0.008 ∼ 0.8 Hz), and smoothing with a full width at half maximum of 3 mm. Nuisance variables, which include six head movement artifacts and the signal from white matter and CSF, were removed using a general linear model. Then we extracted and averaged the values within each segmented ROI. The correlation coefficients between the ROIs were then calculated and used for subsequent analyses.

### Behavior index assessment

#### Goal-directed behavior index

The goal-directed behavior index was classified into two analysis methods, manual and deep learning-based. These behavioral indices are presented in Table S1.

#### Manual analysis

For the goal-directed behavior, we investigated the following behaviors: 1) approach hands to goal [number (n)], 2) bring the goal to the mouth [number (n)], and 3) food peeling [number (n)]. For the ‘approach hands to goal,’ we counted the number of the actual approach to food, spout, or non-food object. The monkey’s act of bringing the goal to their mouth was counted as the index of ‘bring the goal to the mouth.’ Lastly, the number of ‘food peeling’ was defined as when the monkeys peel the food to eat. All behaviors were quantified when the monkey showed low locomotion (body part velocity < 6 px/s). For each behavior, if each low locomotion with the previous 3 s contains the specific behavior, we considered as the low locomotion with respect to that behavior.

#### Deep learning-based analysis

In addition to manual analysis, deep learning-based analysis was used by DeepLabCut. We assessed the three behavioral indices: tray approach [number (n)], bout [number (n)], and duration in food zone [duration (s)].

Throughout the analysis, we utilized several preprocessing methods to improve the accuracy of the results. First, the predicted coordinates of body parts with a likelihood < 0.9 were removed, and those missing values were linearly interpolated using predictions of likelihood > 0.9. Before every computation, this interpolation was conducted. Second, we assumed that the same event (i.e., the hand entering the tray) is not likely to happen multiple times within some short time interval (0.2 s). Hence, even if it is predicted that such an event happened multiple times within 0.2 s, it is reasonable to count only one. This minimum interval constraint is applied while counting the events of a hand entering the tray zone or the mouth zone. Third, before computing the indices related to low locomotion, we averaged over an interval of time the variations of the coordinates; specifically, we computed the moving average of coordinates with a window of 15 s. Those preprocessing methods all contributed to alleviating the effects of inaccurate predictions.

We defined the ‘tray approach’ as the number of moments at which a monkey’s hand entered a tray zone. The ‘duration in food zone’ was defined as the total length of time a monkey remained in the flood zone. Finally, we recursively defined the behavior ‘bout’ as follows: first. We counted the number of events when a monkey’s hand entered the tray zone and subsequently entered the mouth zone within 3 s. In addition, we added the number of events when a monkey’s hand entered the mouth zone within 1 s from the previous ‘bout;’ this defines the behavior ‘bout,’ and we counted the total number of this behavior.

#### Intake

Further, we measured the amount intake of palatable (g), unpalatable food (g), and water (ml) which was freely provided.

### Analysis method

#### Manual analysis

During the experimental period, the behavior of each monkey was recorded by side, top, and front camera at the same time. A clicker synchronized all cameras as a sign of starting the experiment. With the recorded experimental video, the Noldus Observer XT program was used for analysis, and all behaviors were observed at 0.5×-1× speed, shown in Fig. 1A. All experimental manual analysis was done in a blinded manner without knowledge of what drug (e.g., CNO or vehicle) was injected. Only those who conducted the experiment perceived which was LHA^GABA^ neuron activation or control, and the analysts randomly received experimental videos to analyze. There are four manual analysts, and they conducted manual analysis according to the behavior indices. Two manual analysts were paired for each experimental task for accurate and unbiased analysis and did a double-checking analysis.

#### Deep learning-based analysis

To label and predict the coordinates of each body part of DeepLabCut, we manually extracted 240 characteristic frames to label. The extracted images were transformed into training sets, on which the deep neuronal network was trained. The network consists of ResNet-50 (pre-trained on ImageNet) and deconvolutional layers, whose outputs are score maps representing the soft predictions for the location of each body part. The network was trained through 500,000 iterations, minimizing the cross-entropy of the predicted probability distribution relative to the ground-truth probability distribution. After training the model, the coordinates of body parts were predicted in movie S1.

### Operant conditioning paradigm on a fixed ratio 1 (FR1)

The operant conditioning paradigm was conducted using the Cambridge Neuropsychological Test Battery (CANTAB; Lafayette, IN, USA). The CANTAB touchscreen monitor was attached in front of the test cage during the experiment. We selected Monkey A as an experimental subject, with a moderate temper, a high interest in food, and an inquisitive nature suitable for the experiment. The test cage was a stainless-steel cage (111*51*75 (cm)); the monkey’s hand could reach out of the cage and touch the screen. There were plastic doors on each side of the cage to obscure the view of the outside of the cage and focus solely on the CANTAB apparatus. The pellet door for rewards was located on the right side of the touchscreen, and the time was recorded on the system when the monkeys put their hands in the door. The operant conditioning paradigm was a slightly modified version of that which was previously published *(49)*. We used customized sweet pellets with higher sucrose content for the training and experiment as a palatable food. The composition of the pellets is as follows: Sucrose 386.25 g/kg, dextrose 386.25 g/kg, cellulose 127.4 g/kg, grape flavor 30 g/kg, fat 60 g/kg, tableting aids 10 g/kg, and food dye 0.1 g/kg. The monkey was fed 3 hours before the experiment. On an experimental day, the monkey was injected with vehicle or CNO intramuscularly 30 min before the experiment in her home cage and moved to the experimental cage using a transfer cage. The experimental room was separated from the home cage to encourage the monkey to concentrate well on the task. The LHA^GABA^ neuron activation and control effect was measured on a different day every other week to wash out the previous drug. The training for the initial touch on CANTAB was performed with the same protocol as the test without drug injection once a week for three times.

The task began with a green triangle (object) present in the center of the screen. When the monkey touched it, a reward sound and sweet pellet were given, and the object’s color changed to red for 0.2 s. The inter-trial interval (ITI) was 5 s with a black screen. The green triangle remained on the screen until the monkey touched the triangle. The task operated for 30 min. The monkey could freely move in the test cage. Two wireless cameras (FravelCam) were recorded from the test cage (top view) and above the CANTAB (front view) during the experiment. The number of successful trials and door open without the pellet providing (anticipatory behaviors) were calculated to measure the motivation to pellet. Two latencies were used for analysis: the latency between object presence and touching the object (object-touch time) and between touching the object and opening the pellet door (touch-door open time). For calculating the touch-door open time, the first door-open time was followed right after the object touch was used, and the rest of the door-open attempts were ignored.

### Statistical analysis and graph

All the statistical analyses and graphs were performed using Python. For the linear regression in fig. S4, we used GraphPad Prism v.9.3.1 for the analysis. In Fig. 1, D and E, we conducted the paired t-tests to compare each session and normalized value between pre and control, control and activation, and pre and activation. All the individual statistics data are shown in fig. S2 and S3. Further, in Fig. 2, S2, S5, S6, and S7, we conducted the paired t-tests (two-tailed) to compare each index between LHA^GABA^ neuron activation monkeys and control monkeys. Unless zero is contained in the values of the index of control monkeys, we computed the rate of change that occurred by LHA^GABA^ neuron activation; to determine whether those rates are significantly different from zero, we conducted the one-sample t-test for those rates. Furthermore, we compared the total count of successful trials and anticipatory behaviors after the LHA^GABA^ neuron activation and control by bootstrapping, respectively (Fig. 2, K and L). Specifically, we sampled random time windows of equal length of total duration with replacement and compared the number of successful trials between the two conditions within the sampled window. Then the process was repeated 500,000 times. For the statistical analysis of the comparisons between LHA^GABA^ neuron activation and control FC changes within whole brain compartments (Fig. 3), we used the chi-square test. To perform the test, we calculated the frequencies of FC changes that fell above or below the 95% percentile of the distribution of control FC changes. The p-values were the proportion of the difference between the two distributions over zero. Each mean value was presented as mean + SEM. *P*-values < 0.05 were considered significant (*P* < 0.05 (*\**), *P* < 0.01(*\*\**), *P* < 0.001 (\*\**\**), *P* < 0.0001(*\*\*\*\**)). Refer to the Code Availability for the code used for the statistical analysis.

## Acknowledgments

We thank Hyoung F. Kim at Seoul National University, South Korea, for the advice on interpreting our study.

## Author contributions

Conceptualization: LJH, HY, YGK, IB, EB

Methodology: LJH, HY, YGK, IB, YHL, JW, YJ, JP, CJ, KK, JM, YS, JP, EB

Investigation: LJH, IB, HY, YGK, YHL, YJ, JM, YS, JP, KRN

Visualization: LJH, IB, SS, SMY, KRN, JP, EB

Funding acquisition: HC, YL, JC, JYC

Project administration: LJH, SP, WSC, KSL

Supervision: HJC, YL, JC, JYC

Writing – original draft: LJH, HY, YGK, IB, EB

Writing – review & editing: YHL, JC, JW, YJ, JM, YS, JP, KRN, SS, SMY

## Data and materials availability

See https://github.com/Gavroche11/Monkey_chemogenetics for the code related to Fig. 2, and Figs. S5-S9. All other data are available in the main text or the supplementary materials.

## References and Notes

1. A. I. Heriseanu, P. Hay, L. Corbit, S. Touyz, Relating goal-directed behaviour to grazing in persons with obesity with and without eating disorder features. Journal of Eating Disorders 8, 1–14 (2020).

2. A. Noritake, K. Nakamura, Rewarding-unrewarding prediction signals under a bivalent context in the primate lateral hypothalamus. Scientific Reports 13, 5926 (2023).

3. G. D. Stuber, R. A. Wise, Lateral hypothalamic circuits for feeding and reward. Nature neuroscience 19, 198–205 (2016).

4. Alexander C. Whiting, Elizabeth F. Sutton, Corey T. Walker, Jakub Godzik, Joshua S. Catapano, Michael Y. Oh, Nestor D. Tomycz, Eric Ravaussin, Donald M. Whiting, Deep brain stimulation of the hypothalamus leads to increased metabolic rate in refractory obesity. World neurosurgery 121, e867–e874 (2019).

5. A. Noritake, K. Nakamura, Encoding prediction signals during appetitive and aversive Pavlovian conditioning in the primate lateral hypothalamus. Journal of neurophysiology 121, 396–417 (2019).

6. J. H. Jennings, R. L. Ung, S. L. Resendez, A. M. Stamatakis, J. G. Taylor, J. Huang, K. Veleta, P. A. Kantak, M. Aita, K. Shilling-Scrivo, C. Ramakrishnan, K. Deisseroth, S. Otte, G. D. Stuber, Visualizing hypothalamic network dynamics for appetitive and consummatory behaviors. Cell 160, 516–527 (2015).

7. E. Qualls-Creekmore, S. Yu, M. Francois, J. Hoang, C. Huesing, A. Bruce-Keller, D. Burk, H. R. Berthoud, C. D. Morrison, H. Münzberg, Galanin-expressing GABA neurons in the lateral hypothalamus modulate food reward and noncompulsive locomotion. Journal of Neuroscience 37, 6053–6065 (2017).

8. E. H. Nieh, C. M. Vander Weele, G. A. Matthews, K. N. Presbrey, R. Wichmann, C. A. Leppla, E. M. Izadmehr, K. M. Tye, Inhibitory input from the lateral hypothalamus to the ventral tegmental area disinhibits dopamine neurons and promotes behavioral activation. Neuron 90, 1286–1298 (2016).

9. J. N. Siemian, M. A. Arenivar, S. Sarsfield, C. B. Borja, C. N. Russell, Y. Aponte, Lateral hypothalamic LEPR neurons drive appetitive but not consummatory behaviors. Cell reports 36, 109615 (2021).

10. Y. H. Lee, Y. B. Kim, K. S. Kim, M. Jang, H. Y. Song, S. H. Jung, D. S. Ha, J. S. Park, J. Lee, K. M. Kim, D. H. Cheon, I. Baek, M. G. Shin, E. J. Lee, S. J. Kim, H. J. Choi, Lateral hypothalamic leptin receptor neurons drive hunger-gated food-seeking and consummatory behaviours in male mice. Nature Communications 14, 1486 (2023).

11. E. H. Nieh, G. A. Matthews, S. A. Allsop, K. N. Presbrey, C. A. Leppla, R. Wichmann, R. Neve, C. P. Wildes, K. M. Tye, Decoding neural circuits that control compulsive sucrose seeking. Cell 160, 528–541 (2015).

12. M. Navarro, J. J. Olney, N. W. Burnham, C. M. Mazzone, E. G. Lowery-Gionta, K. E. Pleil, T. L. Kash, T. E. Thiele, Lateral hypothalamus GABAergic neurons modulate consummatory behaviors regardless of the caloric content or biological relevance of the consumed stimuli. Neuropsychopharmacology 41, 1505–1512 (2016).

13. V. A. de Vrind, A. Rozeboom, I. G. Wolterink-Donselaar, M. C. Luijendijk-Berg, R. A. Adan, Effects of GABA and leptin receptor-expressing neurons in the lateral hypothalamus on feeding, locomotion, and thermogenesis. Obesity 27, 1123–1132 (2019).

14. J. E. Park, A. C. Silva, Generation of genetically engineered non-human primate models of brain function and neurological disorders. American journal of primatology 81, e22931 (2019).

15. G. Paxinos, X.-F. Huang, A. W. Toga, The rhesus monkey brain in stereotaxic coordinates. (2000).

16. S. Zhang, W. Wang, S. Zhornitsky, C.-s. R. Li, Resting state functional connectivity of the lateral and medial hypothalamus in cocaine dependence: an exploratory study. Frontiers in Psychiatry 9, 344 (2018).

17. S. M. Sternson, D. Bleakman, Chemogenetics: drug-controlled gene therapies for neural circuit disorders. Cell & gene therapy insights 6, 1079 (2020).

18. P. H. Roseboom, S. A. L. Mueller, J. A. Oler, A. S. Fox, M. K. Riedel, V. R. Elam, M. E. Olsen, J. L. Gomez, M. A. Boehm, A. H. DiFilippo, B. T. Christian, M. Michaelides, N. H. Kalin, Evidence in primates supporting the use of chemogenetics for the treatment of human refractory neuropsychiatric disorders. Molecular Therapy 29, 3484–3497 (2021).

19. M. A. Eldridge, W. Lerchner, R. C. Saunders, H. Kaneko, K. W. Krausz, F. J. Gonzalez, B. Ji, M. Higuchi, T. Minamimoto, B. J. Richmond, Chemogenetic disconnection of monkey orbitofrontal and rhinal cortex reversibly disrupts reward value. Nature neuroscience 19, 37–39 (2016).

20. Y. Nagai, E. Kikuchi, W. Lerchner, K. I. Inoue, B. Ji, M. A. Eldridge, H. Kaneko, Y. Kimura, A. Oh-Nishi, Y. Hori, Y. Kato, T. Hirabayashi, A. Fujimoto, K. Kumata, M. R. Zhang, I. Aoki, T. Suhara, M. Higuchi, M. Takada, B. J. Richmond, T. Minamimoto,, PET imaging-guided chemogenetic silencing reveals a critical role of primate rostromedial caudate in reward evaluation. Nature communications 7, 13605 (2016).

21. M. Oguchi, S. Tanaka, X. Pan, T. Kikusui, K. Moriya-Ito, S. Kato, K. Kobayashi, M. Sakagami, Chemogenetic inactivation reveals the inhibitory control function of the prefronto-striatal pathway in the macaque brain. Communications Biology 4, 1088 (2021).

22. K. Oyama, Y. Hori, K. Mimura, Y. Nagai, M. A. G. Eldridge, R. C. Saunders, N. Miyakawa, T. Hirabayashi, Y. Hori, K. I. Inoue, T. Suhara, M. Takada, M. Higuchi, B. J. Richmond, T. Minamimoto, Chemogenetic disconnection between the orbitofrontal cortex and the rostromedial caudate nucleus disrupts motivational control of goal-directed action. Journal of Neuroscience 42, 6267–6275 (2022).

23. M. Kinoshita, R. Kato, K. Isa, K. Kobayashi, K. Kobayashi, H. Onoe, T. Isa, Dissecting the circuit for blindsight to reveal the critical role of pulvinar and superior colliculus. Nature communications 10, 135 (2019).

24. T. Hayashi, R. Akikawa, K. Kawasaki, J. Egawa, T. Minamimoto, K. Kobayashi, S. Kato, Y. Hori, Y. Nagai, A. Iijima, T. Someya, I. Hasegawa, Macaques exhibit implicit gaze bias anticipating others’ false-belief-driven actions via medial prefrontal cortex. Cell Reports 30, 4433–4444. e4435 (2020).

25. P. Vancraeyenest, J. T. Arsenault, X. Li, Q. Zhu, K. Kobayashi, K. Isa, T. Isa, W. Vanduffel, Selective mesoaccumbal pathway inactivation affects motivation but not reinforcement-based learning in macaques. Neuron 108, 568–581. e6 (2020).

26. D. Jeurissen, S. Shushruth, Y. El-Shamayleh, G. D. Horwitz, M. N. Shadlen, Deficits in decision-making induced by parietal cortex inactivation are compensated at two timescales. Neuron 110, 1924-1931. e1925 (2022).

27. T. Tohyama, M. Kinoshita, K. Kobayashi, K. Isa, D. Watanabe, K. Kobayashi, M. Liu, T. Isa, Contribution of propriospinal neurons to recovery of hand dexterity after corticospinal tract lesions in monkeys. Proceedings of the National Academy of Sciences 114, 604–609 (2017).

28. M. Kinoshita, R. Matsui, S. Kato, T. Hasegawa, H. Kasahara, K. Isa, A. Watakabe, T. Yamamori, Y. Nishimura, B. Alstermark, D. Watanabe, K. Kobayashi, T. Isa, Genetic dissection of the circuit for hand dexterity in primates. Nature 487, 235–238 (2012).

29. T. Isa, Double viral vector intersectional approaches for pathway-selective manipulation of motor functions and compensatory mechanisms. Experimental Neurology 349, 113959 (2022).

30. T. Hasegawa, S. Chiken, K. Kobayashi, A. Nambu, Subthalamic nucleus stabilizes movements by reducing neural spike variability in monkey basal ganglia. Nature Communications 13, 2233 (2022).

31. K. Oyama, Y. Hori, Y. Nagai, N. Miyakawa, K. Mimura, T. Hirabayashi, K. I. Inoue, T. Suhara, M. Takada, M. Higuchi, T. Minamimoto, Chemogenetic dissection of the primate prefronto-subcortical pathways for working memory and decisionmaking. Science Advances 7, eabg4246 (2021).

32. N. A. Upright, M. G. Baxter, Prefrontal cortex and cognitive aging in macaque monkeys. American Journal of Primatology 83, e23250 (2021).

33. N. A. Upright, M. G. Baxter, Effect of chemogenetic actuator drugs on prefrontal cortex-dependent working memory in nonhuman primates. Neuropsychopharmacology 45, 1793–1798 (2020).

34. K. Oyama, Y. Hori, Y. Nagai, N. Miyakawa, K. Mimura, T. Hirabayashi, K. I. Inoue, M. Takada, M. Higuchi, T. Minamimoto, Chronic behavioral manipulation via orally delivered chemogenetic actuator in macaques. Journal of Neuroscience 42, 2552– 2561 (2022).

35. M. A. Eldridge, M. C. Smith, S. H. Oppler, J. E. Pearl, J. Y. Shim, W. Lerchner, B. J. Richmond, Unilateral caudate inactivation increases motor impulsivity in rhesus monkeys. Current Research in Neurobiology, 100085 (2023).

36. T. Ninomiya, A. Noritake, K. Kobayashi, M. Isoda, A causal role for frontal corticocortical coordination in social action monitoring. Nature communications 11, 5233 (2020).

37. J. Raper, L. Murphy, R. Richardson, Z. Romm, Z. Kovacs-Balint, C. Payne, A. Galvan, Chemogenetic inhibition of the amygdala modulates emotional behavior expression in infant rhesus monkeys. Eneuro 6, ENEURO.0360-19.2019 (2019).

38. A. Fujimoto, C. Elorette, J. M. Fredericks, S. H. Fujimoto, L. Fleysher, P. H. Rudebeck, B. E. Russ, Resting-state fMRI-based screening of deschloroclozapine in rhesus macaques predicts dosage-dependent behavioral effects. Journal of Neuroscience 42, 5705–5716 (2022).

39. Z. Zhang, L. Shan, Y. Wang, W. Li, M. Jiang, F. Liang, S. Feng, Z. Lu, H. Wang, J. Dai, Primate preoptic neurons drive hypothermia and cold defense. The Innovation 4, 100358 (2023).

40. J. Raper, M. A. G. Eldridge, S. M. Sternson, J. Y. Shim, G. P. Fomani, B. J. Richmond, T. Wichmann, A. Galvan, Characterization of Ultrapotent Chemogenetic Ligands for Research Applications in Nonhuman Primates. ACS Chemical Neuroscience 13, 3118–3125 (2022).

41. K. Mimura, Y. Nagai, K. I. Inoue, J. Matsumoto, Y. Hori, C. Sato, K. Kimura, T. Okauchi, T. Hirabayashi, H. Nishijo, N. Yahata, M. Takada, T. Suhara, M. Higuchi, T. Minamimoto, Chemogenetic activation of nigrostriatal dopamine neurons in freely moving common marmosets. Iscience 24, 103066 (2021).

42. K. Coleman, L. A. Tully, J. L. McMillan, Temperament correlates with training success in adult rhesus macaques. American Journal of Primatology: Official Journal of the American Society of Primatologists 65, 63–71 (2005).

43. D. Weatherall, The Weatherall report on the use of non-human primates in research. London: The Royal Society, 1–145 (2006).

44. N. Percie du Sert, V. Hurst, A. Ahluwalia, S. Alam, M. T. Avey, M. Baker, William J. Browne, Alejandra Clark Innes,C. Cuthill, Ulrich Dirnagl, Michael Emerson, Paul Garner, Stephen T. Holgate, David W. Howells, NatashaA. Karp Stanley, E. Lazic, Katie Lidster, Catriona J. Mac Callum, Malcolm Macleod, Esther J. Pearl, Ole H. Petersen, Frances Rawle, Penny Reynolds, Kieron Rooney, EmilyS. Sena, ShaiD. Silberberg, Thomas Steckler, Hanno Würbel, The ARRIVE guidelines 2.0: Updated guidelines for reporting animal research. Journal of Cerebral Blood Flow & Metabolism 40, 1769–1777 (2020).

45. B. S. Moon, J. H. Park, H. J. Lee, B. C. Lee, S. E. Kim, Routine production of [18F] Flumazenil from iodonium tosylate using a sample pretreatment method: a 2.5-year production report. Molecular imaging and biology 16, 619–625 (2014).

46. Odano, I., Halldin, C., Karlsson, P., Varrone, A., Airaksinen, A. J., Krasikova, R. N., & Farde, L., [18F] Flumazenil binding to central benzodiazepine receptor studies by PET:–Quantitative analysis and comparisons with [11C] flumazenil–. Neuroimage 45, 891–902 (2009).

47. Hyeon-Gu Yeo, Youngjeon Lee, Chang-Yeop Jeon, Kang-Jin Jeong, Yeung Bae Jin, Philyong Kang, Sun-Uk Kim, Ji-Su Kim, Jae-Won Huh, Young-Hyun Kim, Bo-Woong Sim, Bong-Seok Song, Young-Ho Park, Yonggeun Hong, Sang-Rae Lee, Kyu-Tae Chang, Characterization of cerebral damage in a monkey model of Alzheimer’s disease induced by intracerebroventricular injection of streptozotocin. Journal of Alzheimer’s Disease 46, 989–1005 (2015).

48. Colin Reveley, Audr unas Gruslys, Frank Q. Ye, Daniel Glen, Jason Samaha, Brian E. Russ, Ziad Saad, Anil K. Seth, David A. Leopold, Kadharbatcha S. Saleem, Threedimensional digital template atlas of the macaque brain. Cerebral cortex 27, 4463– 4477 (2017).

49. Nicole R Zürcher, Jesse S Rodriguez, Sue L Jenkins, Kate Keenan, Thad Q Bartlett, Thomas J McDonald, Peter W Nathanielsz, Mark J Nijland, Performance of juvenile baboons on neuropsychological tests assessing associative learning, motivation and attention. Journal of Neuroscience Methods 188, 219–225 (2010).

