## Supplementary materials for "Hypothalamic neuronal activation in primates drives naturalistic goal-directed eating behavior"

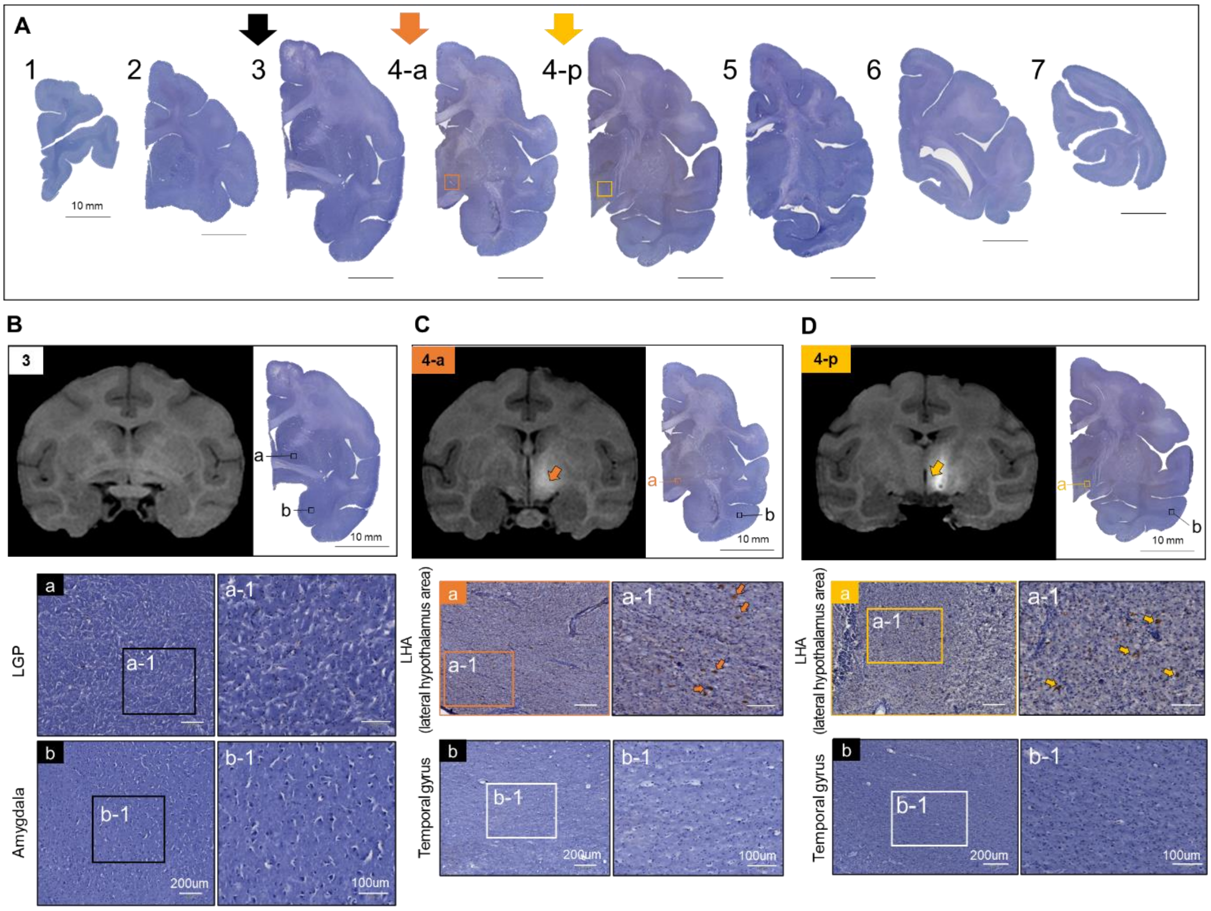


**Fig. S1. A pilot study for virus validation through IHC of Monkey D.** (**A**) The serial samples of brain tissues of Monkey D. a; anterior, p; posterior. Scale bar: 10 mm (**B**) The anti RFP-labeled neurons in slide of 3. a; LGP (lateral globus pallidus), b; Amygdala. (**C**) The anti RFP-labeled neurons in slide of 4-LHA anterior. a; LHA, b; Temporal gyrus. (**D**) The anti RFP-labeled neurons in slide of 4-LHA posterior. a; LHA, b; Temporal gyrus. Scale bar (a, b): 200 µm. Scale bar (a-1, b-1): 100 µm.

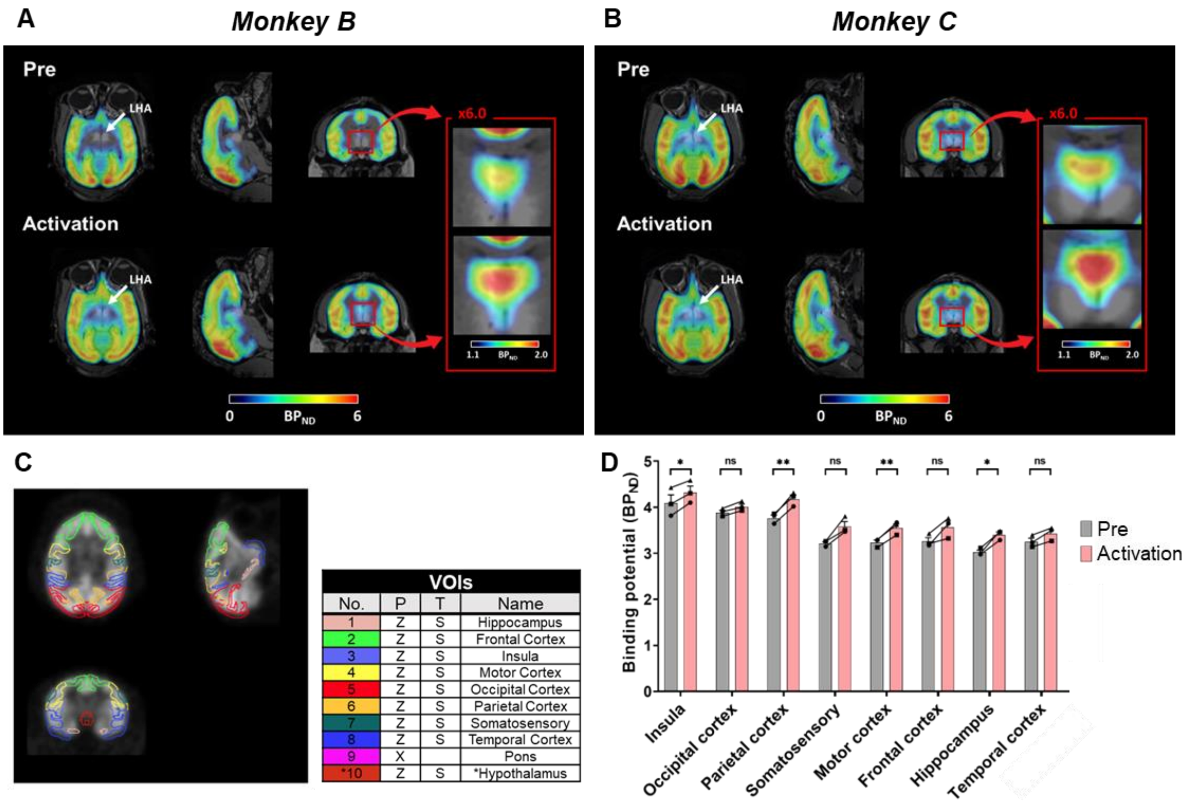


**E**

| | Pre | | | Activation | | | *p Value* |
|---|---|---|---|---|---|---|---|
| | Monkey A | Monkey B | Monkey C | Monkey A | Monkey B | Monkey C | |
| Insula | 3.806711 | 4.058672 | 4.416099 | 4.089927 | 4.293168 | 4.573934 | 0.025272 |
| Occipital cortex | 3.895562 | 3.793135 | 3.966416 | 3.987954 | 3.916668 | 4.123727 | 0.021958 |
| Parietal cortex | 3.633616 | 3.847714 | 3.814088 | 4.008087 | 4.223992 | 4.305283 | 0.008587 |
| Somatosensory | 3.136697 | 3.251425 | 3.254826 | 3.461439 | 3.498108 | 3.793962 | 0.051506 |
| Motor cortex | 3.279221 | 3.120659 | 3.292565 | 3.65957 | 3.389008 | 3.613994 | 0.009858 |
| Frontal cortex | 3.164462 | 3.201672 | 3.419145 | 3.635009 | 3.313819 | 3.754117 | 0.099545 |
| Hippocampus | 2.970252 | 3.107609 | 2.997379 | 3.278594 | 3.448391 | 3.476932 | 0.018925 |
| Temporal cortex | 3.228004 | 3.132944 | 3.39056 | 3.492037 | 3.259393 | 3.541546 | 0.05093 |
| Hypothalamus | 1.019138 | 1.028757 | 1.087872 | 1.19506 | 1.037107 | 1.157145 | 0.226519 |

**Fig. S2. Supplementary data of PET-MR.** (**A**) The comparison of GABA PET-MR images on Monkey B. (**B**) The comparison of GABA PET-MR images on Monkey C. (**C**) Definition of volumes of interests in GABA PET. (**D**) Comparison of binding potential values at pre and $LHA^{GABA}$ neuron activation. (**E**) The individual data of (D). Each value indicated the individual binding potential value of pre, activation, and *p*-value. *p*-value; * $p < 0.05$, ** $p < 0.01$.

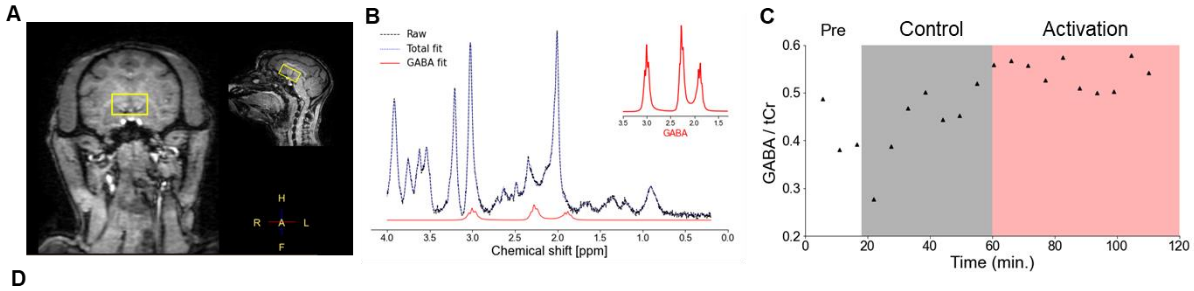


D

| | Pre | | Control | | Activation | | Pre vs. Activation | | Control vs. Activation | |
|---|---|---|---|---|---|---|---|---|---|---|
| | Mean | SD | Mean | SD | Mean | SD | % Change | *p Value* | % Change | *p Value* |
| **Monkey A** | 0.35 | 0.06 | 0.41 | 0.06 | 0.48 | 0.07 | 36.9 | | 17.4 | |
| **Monkey B** | 0.33 | 0.08 | 0.35 | 0.09 | 0.40 | 0.04 | 19.8 | | 15.0 | |
| **Monkey C** | 0.42 | 0.03 | 0.44 | 0.05 | 0.54 | 0.07 | 29.0 | | 24.4 | |
| **All Monkeys** | 0.37 | 0.07 | 0.40 | 0.08 | 0.47 | 0.08 | 28.7 | 0.041 | 19.3 | 0.034 |

*** No significance in Pre vs. Control section**
*** % Change = (Activation-Pre or Control)/(Pre or Control) x 100**

**Fig. S3. Supplementary data of MRS.** (**A**) The brain anatomical reference of MRS. (**B**) The fitting data used by model fitting method of sLASER spectra. (**C**) Real-time dynamics change of individual result on Monkey C. (**D**) The individual statistic data of Fig. 1E. Each value indicated the mean value, SD value, Normalized data (% change), and *p*-value.

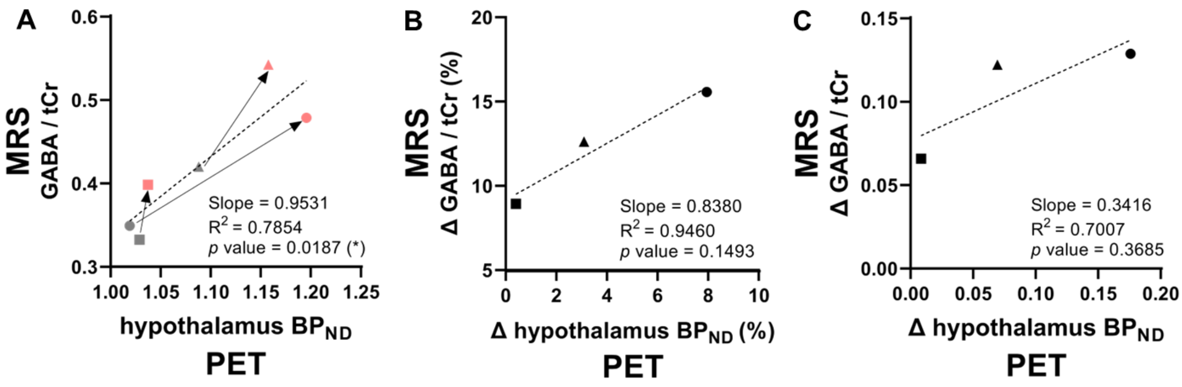


**Fig. S4. The GABA system change value between MRS and PET.** (**A**) Individual value change after the $LHA^{GABA}$ neuron activation through MRS and PET. (**B**) The percentage value differences between MRS (GABA/tCr) and PET (hypothalamus $BP_{ND}$). (**C**) The absolute value differences between MRS (GABA/tCr) and PET (hypothalamus $BP_{ND}$). Throughout the whole figure, the circle, square, and triangle markers correspond to Monkey A, B, and C, respectively. *p*-value; * $p < 0.05$.

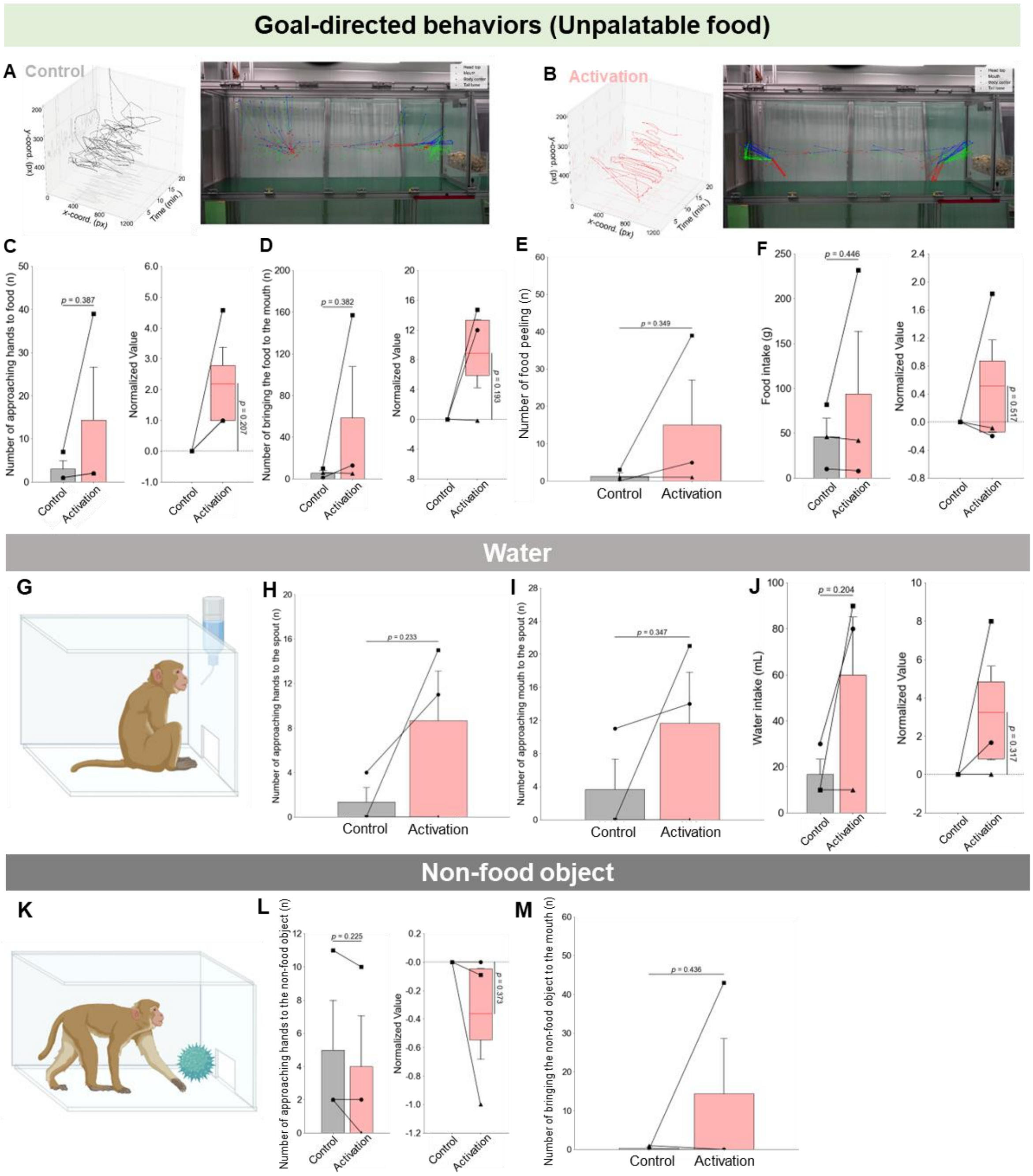


**Fig. S5. No definite change in goal-directed behavior was observed for unpalatable food, water and non-food object after the LHA$^{GABA}$ neuron activation**. (**A**) Behavior temporal dynamics analysis of control. (left) The 3D trajectory of body center (average over an interval time window of 15 sec. length) and (right) the posture of Monkey B (first 5 min.). (**B**) Behavior temporal dynamics analysis of LHA$^{GABA}$ neuron activation. (**C**) The number of approaching hands to food. (**D**) The number of bringing the food to the mouth. (**E**) The number of food peeling. (**F**) The intake of unpalatable food. (**G**) Scheme of water experimental task. (**H**) The number of approaching hands to the spout. (**I**) The number of approaching mouth to the spout. (**J**) The amount of water intake. (**K**) Scheme of non-food object experimental task. (**L**) The number of approaching hands to the non-food object. (**M**) The number of bringing the non-food object to the mouth. The figures in the left side show the values of corresponding indices in Monkey A, B, and C between control and LHA$^{GABA}$ neuron activation (compared by paired t-test). Those in the right side of (C, D, F, J and L) show the rate of change caused by activation (compared with 0 by one-sample t-test). Throughout the whole figure, the circle, square, and triangle markers correspond to Monkey A, B, and C, respectively.

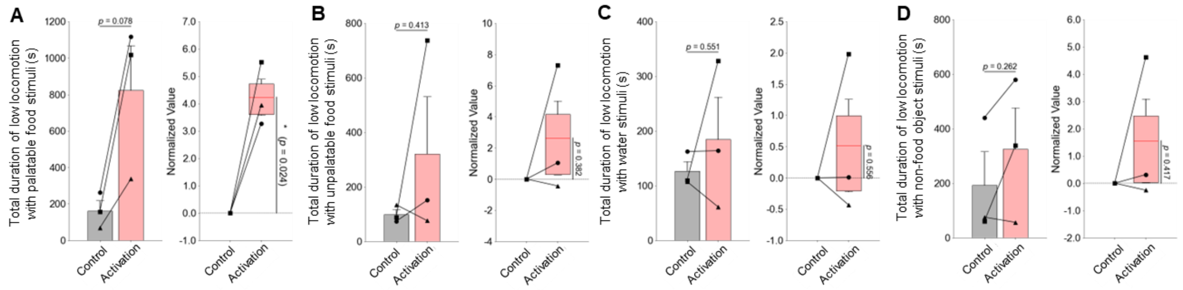


**Fig. S6. The total duration of low locomotion after the LHA$^{GABA}$ neuron activation was significantly increased with palatable stimuli, not with unpalatable, water and non-food object stimuli.** (**A**) The total duration of low locomotion with palatable food stimuli. (**B**) The total duration of low locomotion with unpalatable food stimuli. (**C**) The total duration of low locomotion with water stimuli. (**D**) The total duration of low locomotion with non-food object stimuli. The figures in the left side show the values of corresponding indices in Monkey A, B, and C between control and LHA$^{GABA}$ neuron activation (compared by paired t-test). Those in the right side show the rate of change caused by activation (compared with 0 by one-sample t-test). Throughout the whole figure, the circle, square, and triangle markers correspond to Monkey A, B, and C, respectively. *p*-value; * $p < 0.05$.

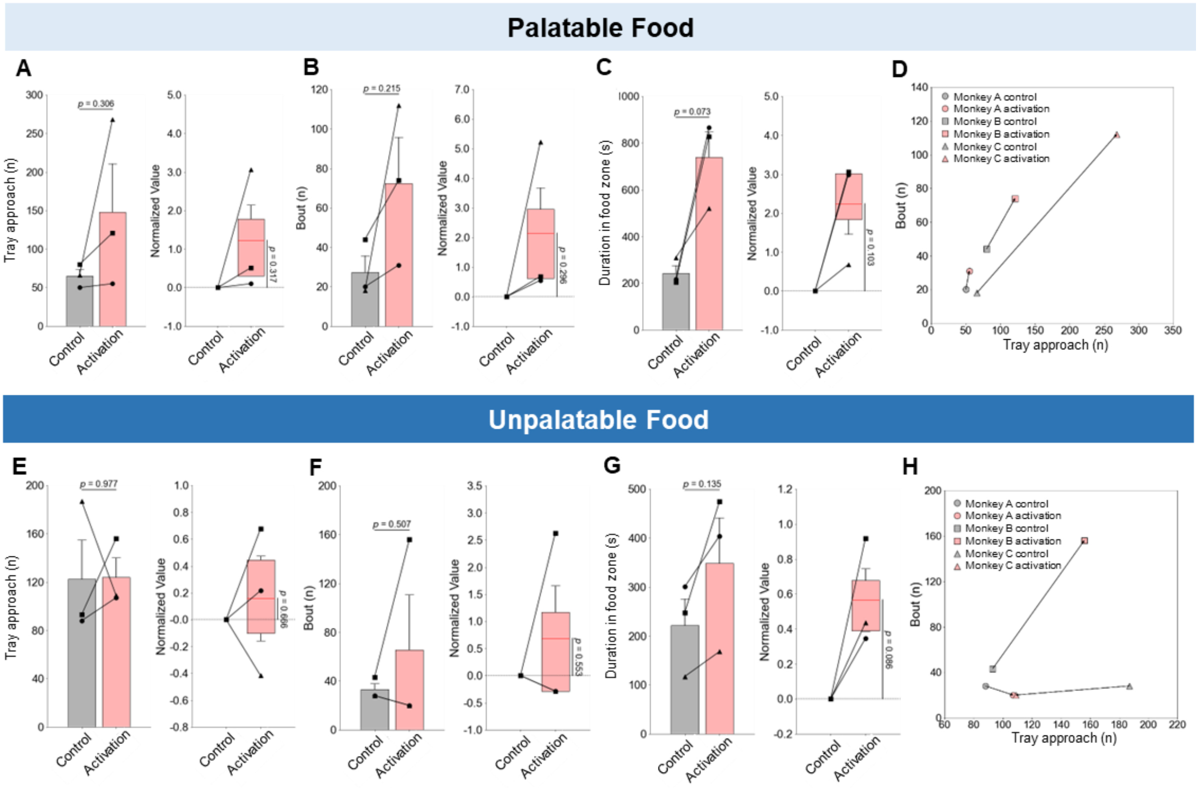


**Fig. S7. Goal-directed eating behavior measured by deep learning-based analysis for palatable and unpalatable food**. (**A-D**) Behavioral indices measured with palatable food stimuli. (**A**) The number of tray approach. (**B**) The number of bouts. (**C**) The duration in food zone. (**D**) The number of bouts plotted over the number of tray approach. (**E-H**) Behavioral indices measured with unpalatable food stimuli. The same behavior indices were plotted as the palatable food stimuli. The figures in the left side of (A-C) and (E-G) show the values of corresponding indices in Monkey A, B, and C between control and $LHA^{GABA}$ neuron activation (compared by paired t-test). Those in the right side show the rate of change caused by activation (compared with 0 by one-sample t-test). Throughout the whole figure, the circle, square, and triangle markers correspond to Monkey A, B, and C, respectively.

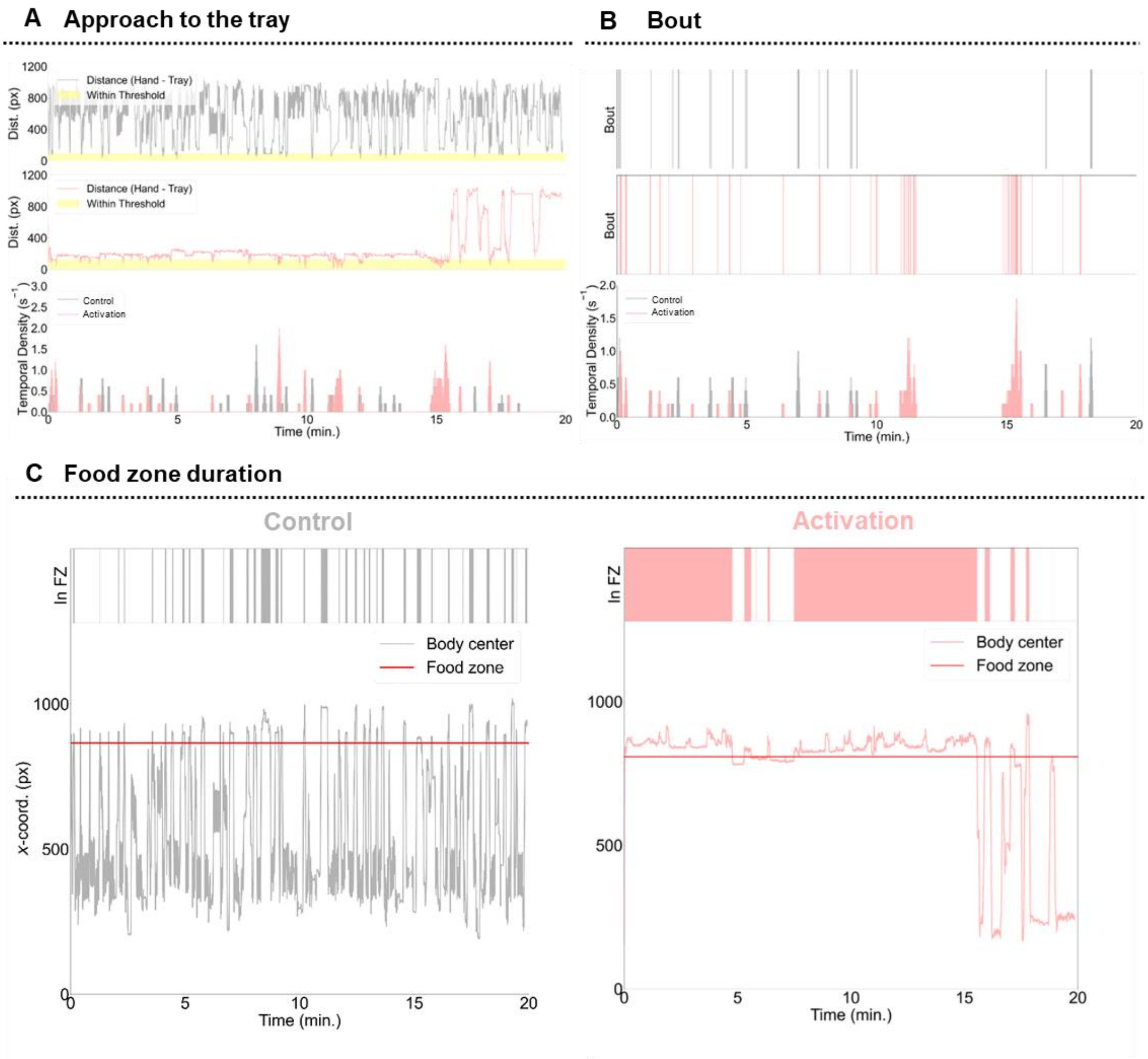


**Fig. S8. Deep learning-based analysis on Monkey B with palatable food stimuli.** (**A**) Distance and temporal density of approach to the tray. Distance between hands and tray over time between (top) control and (middle) $LHA^{GABA}$ neuron activation, where threshold distance is marked as yellow color. (bottom) Temporal densities of tray approach over time between control and $LHA^{GABA}$ neuron activation. (**B**) Frequency and temporal density of bout. Moments of bout over time between (top) control and (middle) $LHA^{GABA}$ neuron activation (bottom) Temporal densities of bout over time between control and $LHA^{GABA}$ neuron activation. (**C**) Duration of food zone time and body center position. (top) Sessions during which the body center of the monkey is located in the food zone and (bottom) *x*-coordinate of body center over time, between (left) control and (right) $LHA^{GABA}$ neuron activation.

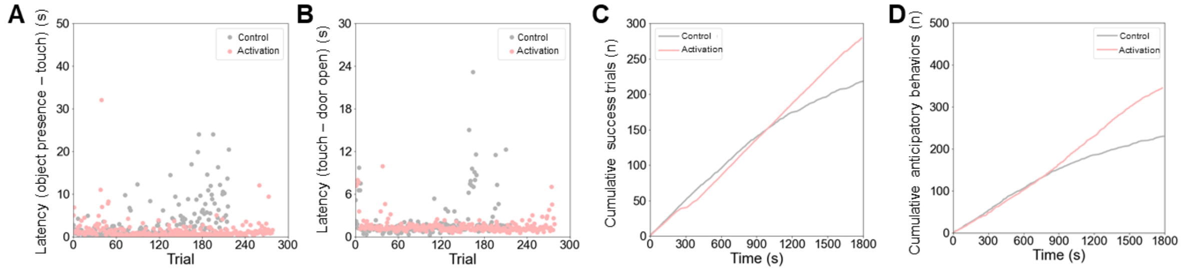


**Fig. S9. Supplementary data of goal-directed motivation.** (**A**) Latency between object presence and object touch for each trial. (**B**) Latency between object touch and door open for each trial. (**C**) Cumulative numbers of success trials over time between control and $LHA^{GABA}$ neuron activation. (**D**) Cumulative numbers of anticipatory behaviors over time between control and $LHA^{GABA}$ neuron activation.

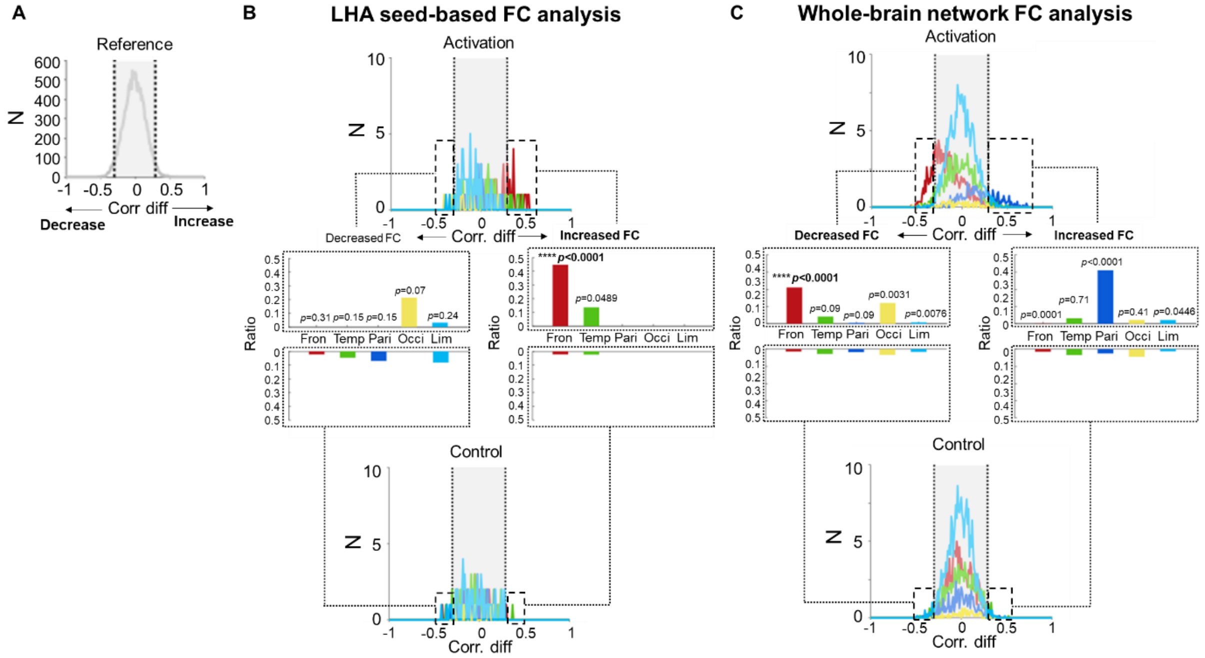


**Fig. S10. LHA seed-based and whole-brain network distribution of FC changes (ΔFCs).** (**A**) A control distribution of ΔFCs (middle, post control – pre control) was constructed, and threshold values (dotted lines, 95%) were obtained (-0.2976 and 0.2889) (**B**) LHA seed-based distributions of ΔFCs were grouped by lobe compartments, with each color representing a different lobe. ΔFCs that deviated from the threshold were counted as significant (shown in the dotted box in the distribution plots). The ratio plots in the middle represents the proportion of significant ΔFCs compared to all possible ΔFCs in the lobe. (**C**) Whole-brain network distributions of ΔFCs (black rectangle in Fig. 3D) were constructed in the same way as in (B). *p*-value; **** p < 0.0001.

| Analysis method | Measured Behavior | Index | Palatable food | Unpalatable food | Water | Non-food object |
|---|---|---|---|---|---|---|
| Manual analysis | Approach hands to goal | Number (n) | ↑ * | ↑ | ↔ | ↔ |
| | Bring the goal to the mouth | Number (n) | ↑ * | ↔ | ↔ | ↔ |
| | Food peeling | Number (n) | ↑ * | ↔ | N.A. | N.A. |
| Deep learning-based analysis | Tray approach | Number (n) | ↑ | ↔ | N.A. | N.A. |
| | Bout | Number (n) | ↑ | ↔ | N.A. | N.A. |
| | Duration in food zone | Duration (s) | ↑ | ↑ | N.A. | N.A. |
| Intake | Food / Water intake | Amount (g / mL) | ↑ | ↔ | ↔ | N.A. |

**Table S1. The behavioral index and summary of goal-directed behavior**. Efficacy summary of goal-directed behavior after the $LHA^{GABA}$ neuron activation. Upward arrow with asterisk (↑*), significant increase compared to control; Upward arrow (↑), consistent increase for all three monkeys compared to control; ↔, no significant or consistent change for all three monkeys compared to control; **N.A**, not applicable. *p*-value; * $p < 0.05$.

**movie S1.**

Goal-directed eating behavior for palatable food ($LHA^{GABA}$ neuron activation)

**movie S2.**

Goal-directed motivation for palatable food using operant conditioning paradigm ($LHA^{GABA}$ neuron activation)

**movie S3.**

Goal-directed motivation for palatable food using operant conditioning paradigm (control)
